# Gene expression patterns decompose fMRI activation in a sub-region-specific manner in mice after nociceptive stimulation

**DOI:** 10.1101/2025.04.01.646627

**Authors:** Isabel Wank, Anja Amthor, Silke Kreitz, Maximilian Häfele, Andreas Hess

## Abstract

The results of functional magnetic resonance imaging (fMRI) cannot be interpreted directly at the molecular or genetic level. Our integrative workflow aims to add such a new dimension to the functional interpretation using a publicly available mouse brain gene expression database from the Allen Institute for Brain Science (ABA). From an average of 164 in-house measured mouse thermal pain fMRI datasets, we identified the top and bottom 5% of voxels according to their activation probability (AP) in response to warm and hot hind paw stimulation. Here, we investigated whether high AP voxels differ from low AP voxels in terms of gene expression: Analyzing nine core brain regions of the ‘pain’/saliency system, the top (high AP) and bottom (low AP) 5% of voxels showed distinct gene expression profiles. In nearly all regions, only high AP voxels were significantly enriched for gene ontology (GO) terms related to neurotransmitter activity, synaptic structure and neuronal function, while only the dorsal striatum showed GO term enrichment in low AP voxels. Notably, randomly selected voxels showed no significant enrichment, demonstrating the reliability of this approach.

These results highlight the potential of gaining knowledge by integrating gene expression and fMRI data. Despite the limitations of using fixed gene expression data from the ABA cohort, this approach may provide new insights into physiological processes and improve the parcellation and interpretation of imaging data.

## Introduction

Interpreting the biological significance of functional magnetic resonance imaging (fMRI) data in preclinical imaging can be complex and therefore challenging. Although fMRI can detect brain activity and its changes or abnormalities with high sensitivity, it is often difficult to understand what causes these changes or what they imply for such as behavioral tests or, more recently, genetic or omics data. However, omics data often lack 3D spatial resolution. Here, we present a workflow that integrates high-resolution spatial gene expression data from the publicly available Allen Mouse Brain Atlas (ABA) from the Allen Institute for Brain Science with stimulus-evoked fMRI data to enhance data interpretation [1, 2].

Although the genomic DNA is identical across all cells of our body, the actual gene expression (the transcriptome) varies greatly between cells. This variability is influenced by DNA structural re-organizations, such as histone modifications like methylation and acetylation [3] or, even faster by expression of immediate early genes like c-Fos [4, 5]. This can result in (de-)activation of substantial portions of DNA, relocation of enhancer regions and alteration of the transcription factor and repressor landscape, ultimately affecting gene expression.

Differences in the transcriptome contribute to the diversity of cell phenotypes, functions, and morphologies necessary for complex biological processes [6]. However, cells with (slightly) different transcriptomes can still perform the same physiological task due to redundancy in protein function [7].

The brain contains four main cell types: neurons, glial cells, endothelial cells, and pericytes [8]. The glial cells have supportive functions, such as providing nutrients to neurons and building myelin sheaths, but also have immune functions as scavenging microglia [9]. Neurons, the primary signaling cells, are further classified by function (excitatory, inhibitory, and neuromodulatory) as well as morphology. They are organized into circuits that span multiple brain regions and can exhibit different behaviors depending on their location. Certain neurons respond specifically to sensory stimulation, and there is even a small subset of neurons that respond dominantly to nociceptive stimuli [11–14], activates a broad network of regions involved in sensory, emotional, motor, cognitive, and autonomic processing that can be mapped using fMRI [15–17]. Despite the relatively low spatial resolution used in mouse studies (here 234 µm x 234 µm x 500 µm), the small changes in brain hemodynamics can still be traced to the individual voxel level, making it possible to identify also the regions of the mouse brain that process noxious stimuli [16, 18].

The Allen Institute for Brain Science provides in situ hybridization (ISH) brain gene expression data for 19,422 genes derived from 8-week-old male C57BL/6J mice [1, 2], mapped in a 200 µm x 200 µm x 200 µm grid together with a corresponding brain atlas for identification of brain regions [19]. With an average cortical neuron density in the mouse brain of 92,616 ± 25,000 cells/mm^3^ (mean ± std) [20], such a single voxel contains approximately 740 neurons.

This study aimed to determine whether voxels that are most likely to be activated by noxious thermal stimuli show distinct gene expression patterns compared to voxels with low activation probability (AP), suggesting a localized higher abundance of genetically defined highly specialized neurons in these voxels.

To draw conclusions about the differential gene expression patterns between high and low AP voxels, the gene set enrichment analysis (GSEA) allows a functional characterization of the extensive gene expression data. We used GSEA to investigate spatial gene expression patterns for biological functions and cellular components, helping us to interpret our imaging data in a more biological context. Finally, but importantly, we attempted to falsify the approach by comparing two randomly selected non-overlapping voxel sets.

## Results and Discussion

### Spatial distribution of activation probability

We selected nine core regions (S1 Table) of the brain’s ‘pain matrix’, a system that detects not only pain, but processes all kinds of highly salient stimuli [11, 21] as regions of interest.

**S1 Table.** Overview over the analyzed brain regions. Top part are the nine core regions, below are three regions analyzed for proof of concepts (activated by thermal stimuli) but not core regions of the brain’s ‘pain’/saliency system.

| Name | Abbreviation | Brain regions contained | Volume in voxels |
| --- | --- | --- | --- |
| Thalamus | TH | thalamus | 2552 |
| Somatosensory Cortex | CTXs | primary and secondary somatosensory cortex | 4828 |
| Cingulate Cortex | CG | cingulate and retrosplenial cortex | 1739 |
| Insular Cortex | INS | anterior and posterior insular cortex | 643 |
| Visual Cortex | VIS | visual cortex | 1667 |
| Hippocampus | HC | dendate gyrus, subiculum and hippocampus | 3924 |
| Dorsal Striatum | STRd | caudoputamen | 3142 |
| Ventral Striatum | STRv | nucleus accumbens, olfactory tubercle, islands of Calleja, fundus of striatum | 1148 |
| Cerebellum | CER | cerebellum | 7900 |
| Motor-related<br>Pons | PONmr | motor-related parts of<br>the pons | 1854 |
| Piriform Cortex | PIR | piriform cortex | 2090 |
| Hypothalamus | HYP | hypothalamus | 1632 |

Brain regions often have a distinct cellular composition, which has led to histological identifiability (cf. e.g. Brodmann areas [22] or Paxinos [23]). In contrast, the cortex is assumed to be relatively similar histologically (6-layer structure). It is noteworthy that there are sharp functional divisions within this histologically homogeneous (sensory) cortex. For example, the auditory cortex shows a clear and sharp boundary to the neighboring somatosensory cortex, which can be revealed by specific GABA receptor activation [24]. Following this idea of spatial functional sub-segregation within the same histologically defined structure (also found in resting state analyses of the (human) brain), we first asked whether voxels with high vs. low fMRI AP to thermal stimuli are heterogeneously distributed, i.e. can be spatially separated. Second, if such a heterogeneous representation exists, we asked if this could be attributed to a differential gene expression within the same brain region. Activation probability maps were derived from the group average of 164 individual mouse fMRI measurements, containing only the significantly activated (binarized) voxels per animal evoked by either warm or hot stimulation temperatures (see Methods for details).

Interestingly, we found a distinct spatial segregation of low and high AP voxels per brain region (S1 Fig). In some cases, such as CTXs, VIS and HC, the distribution pattern correlated well with functional subdivisions. For example, the mouse HC can be functionally divided into dorsal (higher AP), memory-related and ventral (lower AP), emotion/stress-related parts [25]. This led us to ask whether the differences in AP are reflected in underlying gene expression, determining the different physiological functions of these regions.

To assess gene expression differences accordingly, we sorted all voxels for each region according to their AP in descending order and analyzed the gene expression of the top and bottom 5% voxels (S2 Fig), i.e. the 5% voxels with the lowest and highest AP per brain region (for specific voxel numbers see S2 Table in Methods). Fig 1 shows the spatial distribution of the resulting voxels analyzed.

**Fig 1.**
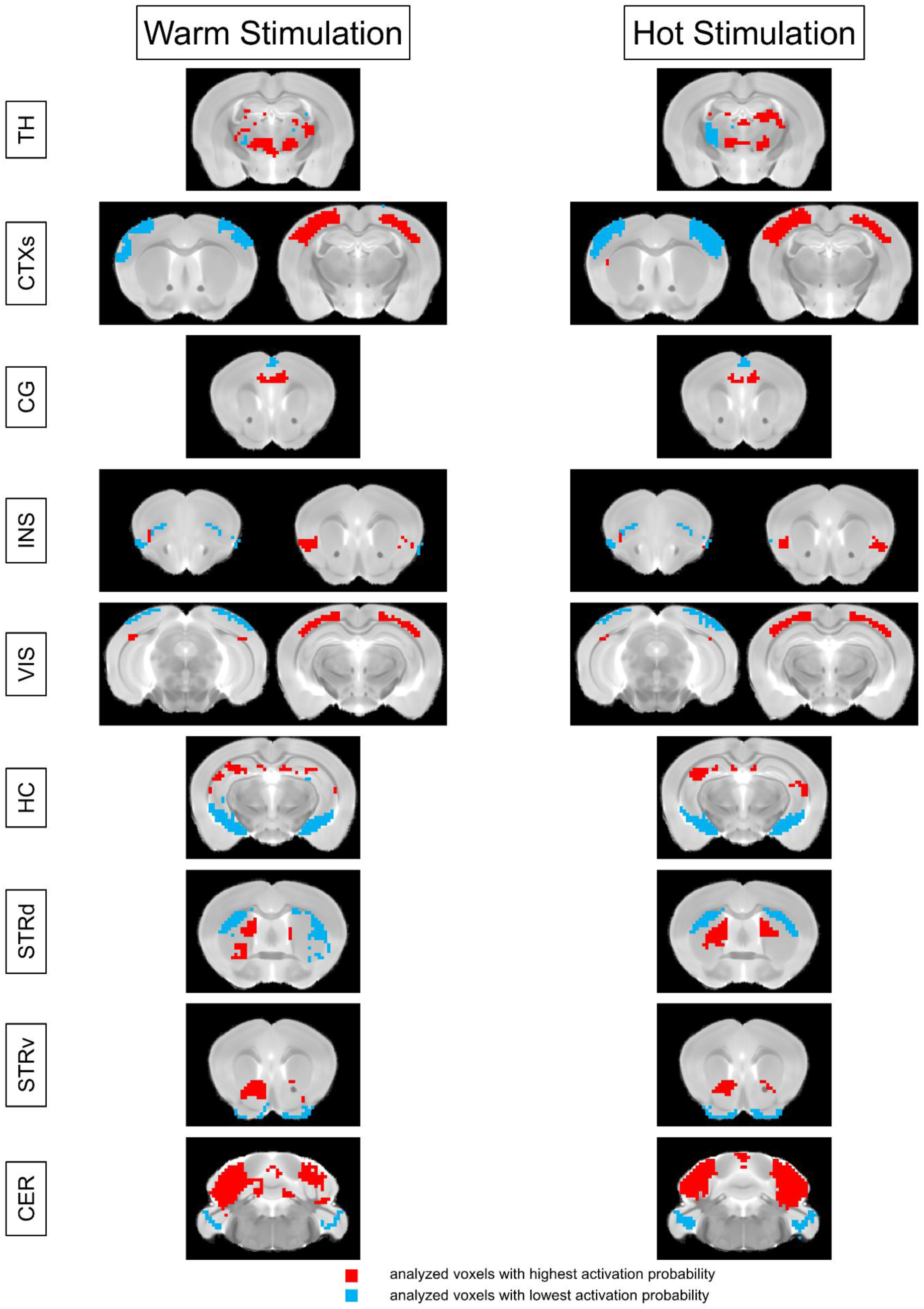
Spatial distribution of the 5% analyzed voxels. Shown are representative slices for the brain regions analyzed by GSEA. The top and bottom 5% analyzed voxels were averaged across all genes to identify the spatial distribution of the voxels selected for analysis. Blue voxels represent the lowest AP, red voxels represent the highest AP.

**S2 Table.** Voxel numbers analyzed using GSEA per brain region. Given is the number of the top and bottom 5% voxels with highest/lowest AP (for thalamus, 128 voxels with the highest and 128 voxels with the lowest probability were analyzed). Abbreviations: TH - thalamus, CTXs - somatosensory cortex (including S2), CG - cingulate and retrosplenial cortex, INS – insular cortex, VIS - visual cortex, HC - hippocampus (including subiculum and dendate gyrus), STRd - dorsal striatum, STRv - ventral striatum (including nucleus accumbens and olfactory tubercle), CER - cerebellum.

| Brain Region | TH | CTXs | CG | INS | VIS | HC | STRd | STRv | CER |
| --- | --- | --- | --- | --- | --- | --- | --- | --- | --- |
| Voxel Number | 128 | 214 | 87 | 32 | 83 | 196 | 157 | 57 | 380 |

In all regions analyzed, the spatial distribution of the voxels with the lowest AP markedly differed from that of voxels with the highest AP. Despite being derived from two different AP maps, the distribution for warm and hot stimulation temperatures was highly consistent, indicating that the underlying spatial activation probability patterns reliably represent similar driving physiological processes (Fig 1 left (warm) vs right (hot)) [26].

Next, the range of AP for the top and bottom 5% voxels (significantly activated by exactly the same stimulus) of each brain region was analyzed (Fig 2). As expected, warm stimulation led to lower AP than hot stimulation in all brain regions analyzed. This observation is consistent with many previous findings (e.g. [17]) that hot stimulation typically elicits stronger and more widespread fMRI activations, likely due to its higher intensity and greater involvement of sensory and nociceptive pathways.

**Fig 2.**
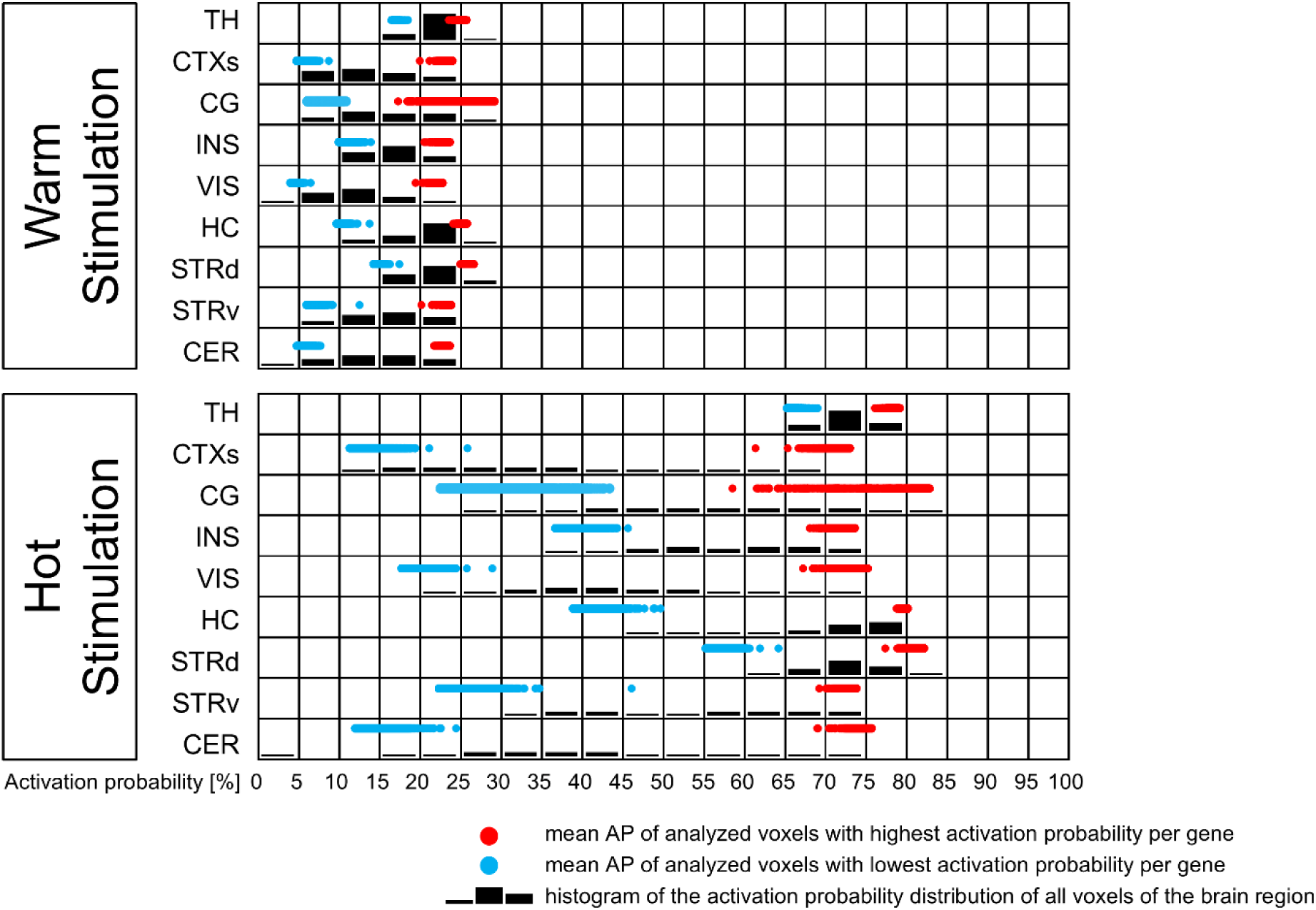
Distribution of AP. The black bars represent a histogram of the frequency of the AP (x-axis) across all voxels of the brain region. Blue and red dots represent the average AP of the top and bottom 5% voxels analyzed for each gene.

Across brain regions, the range of AP between bottom and top 5% voxels was quite heterogeneous. For example, in TH and STRd, the AP values of both bottom and top 5% voxels were relatively high on average, resulting in a smaller range compared to other regions. This implies a less distinct separation between the bottom and top 5% voxels and suggests that even the low AP voxels in TH and STRd still showed frequent responses to stimulation (e.g. more than in CER or VIS). In contrast, CTXs and CER had similarly high AP values for the top 5%, but much lower AP values for the bottom 5% voxels, indicating a greater separation between the bottom and top 5% voxels.

### Selective functional enrichment of gene ontology terms in high AP voxels

There are several ways to functionally characterize large gene expression data sets. Online databases such as PANTHER [27], TOPPGene Suite [28] or DAVID [29] provide functional genomic annotations for lists of genes that differ between the datasets of interest. However, these lists are generated a priori by the researcher and are not weighted by the actual expression data, but all genes are treated equally. Initially, we started our approach with these databases but then switched to gene set enrichment analysis [30]. Like PANTHER, TOPPGene Suite and DAVID, the algorithm used in GSEA calculates whether gene ontology (GO) terms are enriched in given gene expression data. However, unlike the previous databases, GSEA is based on individual replicate expression values (here expression values from many single voxels) and includes a rank-based weighting mechanism to account for real differences in gene expression levels in the compared data sets.

GO terms describe biological processes (**BP**, e.g. ‘dopamine secretion’), cellular components (**CC**, e.g. ‘terminal bouton’) or molecular functions (e.g. ‘ion channel’). Each GO term contains a list of all genes associated with that term. The rank-based statistical analysis calculates whether these genes are overrepresented in the expression data of one group or the other. As a result, the algorithm identifies significantly (FDR-corrected) enriched GO terms in each of the compared data sets, allowing biologically relevant conclusions to be drawn.

Comparing the gene expression data of 19,410 genes for the top and bottom 5% voxels with the highest and lowest AP, GSEA ranks and weights the differential gene expression data and calculates for each GO term whether the expression of the genes contained in the term are enriched in high or low AP voxels. The results include a list of the (significantly) enriched GO terms for both low and high AP voxel sets, the number of genes contained in each GO term gene list, the (normalized) enrichment score, and several statistical measures including the FDR q-value to identify statistically significant enriched GO terms.

S3 Fig summarizes the number of enriched GO terms detected by GSEA for the gene sets ‘biological process’ and ‘cellular component’ for warm and hot stimulation. Overall, drastically more (significantly) enriched GO terms were found in the top 5% voxels (S3 Fig, right side) than in the bottom 5% voxels. Significant enrichment (FDR < 0.05) was found almost exclusively for the high AP voxels, with the notable exception of the dorsal striatum for **CC** and hot stimulation. Significant enrichment was found for **BP** in VIS and STRv for both warm and hot stimulation. For **CC**, significant enrichment for warm stimulation was found for CTXs (minor), VIS, STRv, and CER. For hot stimulation, significant enrichment was found for TH (minor), VIS, and STRv.

Further examination of the GO terms that were significantly enriched (FDR q < 0.05) in the high AP voxels, revealed that many of them were related to neurotransmitters, synaptic and neuronal structure and function, cognition, and perception. It is worth mentioning that many GO terms in low AP voxels were related to immunological processes (but insignificant).

Consequently, we counted all significantly enriched GO terms with association with neurobiological structure and function (**neuro-associated GO terms** (FDR q < 0.05)). Those were mainly enriched in the high AP voxels, underlining the specificity of the approach (Fig 3, right side). The number of neuro-associated GO terms was higher than the expected percentage of neuro-associated GO terms in the respective term list. The main exception was again the STRd, where neuro-associated GO terms were found for low AP voxels for **CC** and hot stimulation. No significant enrichment at all was found in CG, INS, and HC.

**Fig 3.**
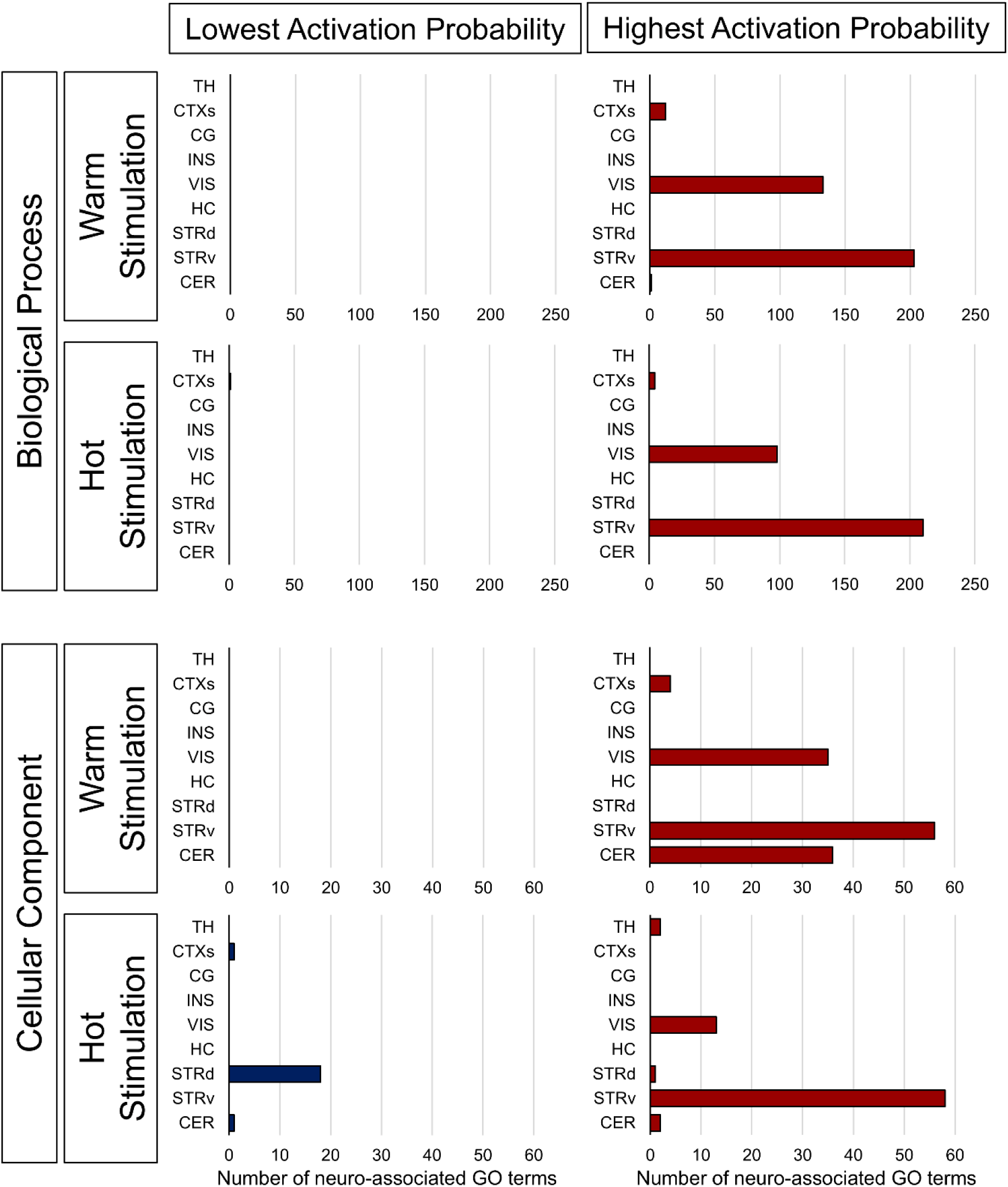
Number of significantly enriched neuro-associated GO terms in the top and bottom 5% voxels. Shown are the number of neuro-associated (GO terms associated with neurons, neurobiological processes or neuronal localizations) GO terms with FDR < 0.05 (usually used in imaging data; blue (bottom 5% voxel set with lowest AP), red (top 5% voxel set with highest AP). For insignificantly enriched GO terms see S3 Fig.

The results suggest that a large range between the mean AP of the top and bottom 5% voxels may promote the enrichment of GO terms, as seen in VIS, STRv, and CER. To test this, we analyzed two additional regions with huge differences between the mean AP of the 5% top and bottom voxels (the motor-related pons (PONmr) and the piriform cortex (PIR)) and one with a notably small difference (hypothalamus (HYP)) (S4A Fig). Again, the regions showed clear spatial separation between low and high AP voxels (S4B,C Fig). However, significant enrichment (FDR < 0.05) was only observed for high AP voxels and warm stimulation in PONmr and HYP for the **CC** gene set (S4D Fig).

Interestingly, small AP differences imply a more homogeneous response across the region. This suggests here less functional differentiation between low and high AP voxels, which could explain the lack of GO term enrichment. This was confirmed in the case of TH, but not for HYP or STRd, where small differences in AP also led to significant enrichment.

We conclude that the difference in AP range between the top and bottom voxels does not influence GO term enrichment.

### Enrichment of neuro-associated GO terms in voxels with high AP

To analyze neuro-association in more detail, we manually curated a neuro-associated subset of GO terms for **BP** and **CC**. Using the Cytoscape ‘Enrichment Map’ plug-in [31], we grouped the neuro-associated **BP** GO terms into functionally related clusters. The algorithm identified 19 distinct clusters covering categories such as neurotransmitter activity, receptor function, synaptic structure and function, neuronal architecture, and higher cognitive functions. These clusters were consolidated into 19 comprehensive GO term groups, which we termed Super-GOs, by aggregating all genes associated with the GO terms within each cluster (see S4 Table). Next, GSEA was performed again using only this Super-GO gene set. Significant enrichment was found for CTXs, INS, VIS, STRv and CER for the high AP voxels, and for STRd and CG only for the low AP voxels (Figs 4 and S5). In the case of VIS and STRv, almost all 19 Super-GOs were significantly enriched. In the case of CTXs (especially for hot stimulation), INS (warm stimulation), and CER (especially for hot stimulation), the enrichment was focused on specific Super-GOs. The enriched Super-GOs for CTXs were acetylcholine, GABA, myelination, dendrite, and axon regeneration. For INS catecholamine, acetylcholine, glutamate, GABA, and cognition and for CER neurotransmitter receptors, receptor activity, action potential, synapse, and neuron/axon.

**Fig 4.**
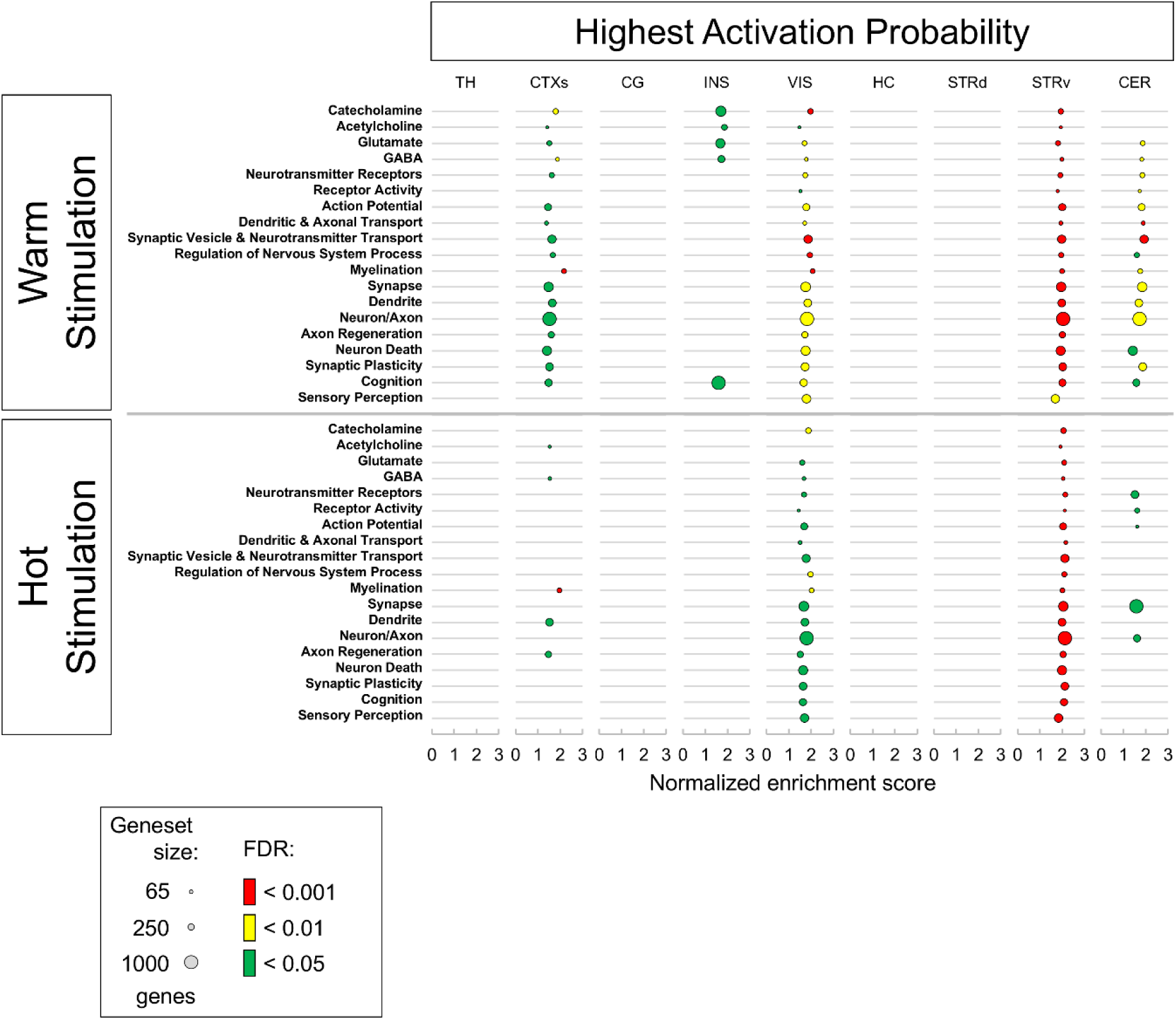
Enrichment of the summarized Super-GO terms for active voxels using the manually curated neuro-associated BP gene list. Bubble size represents the number of genes included in the summarized gene set, color codes the FDR q-value and x-axis shows the normalized enrichment score (NES).

Overall, this suggests that the integration of gene expression patterns (even not from the same animals) and fMRI activation probability can indeed provide new insights and add valuable information to the neurochemical / molecular interpretation of functional MRI data.

Interestingly, one of the two brain regions that always showed significant enrichment with somatosensory thermal stimulation paradigm was the visual cortex. Using somatosensory stimulation paradigms, we often find activation of the visual and auditory cortices, as well as the superior and inferior colliculi. One reason for this activation could be a general increase in the brain’s arousal induced by salient stimuli, which in turn modulates thalamic gating functions [32], leading to broader sensory input forwarding. A second reason could be that although there are clear biochemical boundaries between visual and somatosensory cortex [24], the cortex is known to be not strictly unimodal on a continuous basis [33, 34]: many studies have shown that especially belt regions between visual, auditory, somatosensory and even associative cortex respond to multisensory stimulation (e.g. touch), whereas the core areas seem to be mainly unimodal [35, 36]. Indeed, we found high AP voxels of VIS located at the borders to associative and somatosensory cortex (Figs 1 and S6), while the core area of the visual cortex contains voxels with low AP. Thus, we appear to be comparing unimodal with multimodal ‘voxels’, resulting in the strong enrichment of neuro-associated GO terms in the latter. The somatosensory cortex is highly specialized for processing somatosensory stimuli and is typically more modality-specific, which could be the reason why we find less neuro-enrichment in the somatosensory cortex.

### Validation of analysis specificity using random voxel permutation

To falsify the rationality of the whole analysis approach, we repeated the entire GSEA analysis for the regions CTXs, VIS, STRd, and STRv using random voxels by permuting the previously sorted order (by AP) of the voxels. From this permuted list, Spatial projection demonstrated successful randomization (S7A Fig). The number of overall enriched GO terms was comparable for the two random voxel sets. In contrast to the low and high AP voxels, the random voxels showed no significant enrichment (and consequently no significantly enriched neuro-associated GO terms) (S7B Fig).

### Critiques and Caveats

First, as a general statement, the transcriptome does not necessarily predict the proteome. Consequently, the exact protein composition cannot be directly deduced from gene expression alone.

The key finding of our approach is the enrichment of neuro-associated GO terms in voxels with high AP by thermal stimulation. From a biological point of view, it is difficult to argue for such a significant difference in highly specialized gene expression between low and high AP voxels, since a 200 µm voxel contains many different cell types, and the voxels with lowest and highest AP are sometimes direct neighbors which were thought to have similar cellular composition. But something must distinguish the voxels that respond to a given stimulation from those that do not, and besides differences in connectivity, gene expression patterns are the most likely source.

The gene expression data used in this study were obtained from p56 male C57BL/6J mice from the Allen laboratory. To provide the most reliable cross-section of fMRI response patterns in the general mouse population, we combined fMRI data from male and female C57BL/6N mice and female DBA1 mice (see methods for n-numbers) [37–39]. Therefore, the exact gene expression of our animals used for fMRI may vary. Breeder, age, housing conditions and the fMRI treatment could also influence gene expression [40]. However, as we used average activation probability maps from 164 fMRI measurements, this could be considered broad enough to estimate a generalized conclusions should be drawn with caution for cohorts other than male C57BL/6J mice, which were used for the ABA. However, our results indicate that there is a more specific genetic sub-parcellation beyond strain, gender, breeder, age and laboratory.

In future projects, we plan to incorporate spatial transcriptomics data from our own mice using the MERSCOPE spatial imaging platform’s MERFISH method, which will allow us to directly correlate gene expression and MRI data from the same animal cohort.

Regarding the quality of the gene expression data, which were all derived by ISH staining, all data underwent a rigorous quality assessment at the Allen Institute for Brain Science. However, when comparing ISH images of different genes, some problems were identified, such as slices that clearly did not belong to that gene (no expression in any brain region except in one slice of the cerebellum; Ahr - RP_050407_01_C09 - sagittal), registration errors (detectable due to misalignment of an air bubble artefact in two consecutive slices), artefacts such as enrichment of ISH signal in tissue tears and air bubbles (Bdnf - RP_071204_02_D03 - coronal). Due to the large amount of data, it was not feasible to manually review all 25,520 raw data sets and delete problematic data sets.

The data presented by the Allen Institute may be in coronal and/or sagittal orientation. In case of sagittal data, the dataset consists of the left hemisphere only. We imputed data from the missing hemisphere to make the datasets usable for our pipeline (see Methods), but it should be noted that this is artificial bilaterality and these data cannot be used to infer laterality effects.

To be able to use the ground truth of the ABA gene expression, it was necessary to register our 22-slice fMRI data into the 66-slice ABA space while also adjusting the analyzing only the correspondent gene expression slices, leads to only marginal changes (data not shown). Therefore, we decided to use the registration method to increase the number of voxels analyzed and thus the statistical power of the analysis. We have found that automatic registration approaches such as ANTs [41] place too much emphasis on the outline of the brain. As the GE-EPI fMRI acquisition technique overestimates the thickness of the cortex, the inner brain regions such as the hippocampus and thalamus are shifted inwards and reduced in size in such outline-based registration methods. The resulting registration quality was inadequate, so we opted for the manual landmark-based thinplate registration of 3D Slicer [42]. In general, we achieved good results with about 80-100 individual landmarks (here we used 88). Interpolation steps (especially to create 66 slices from our 22 slices) can introduce data distortion. We found an artificial introduction of negative AP values in voxels at the direct border between brain and background and voxels with an AP less than 1/164 (number of measurements included in the mean fMRI activation probability datasets). Therefore, we restricted the analysis to include only voxels with a minimum AP of 1/164. Nevertheless, a better automatic registration approach should be developed.

GSEA calculates enrichment of gene lists. These lists consist of the GO term and all genes known to be associated with that GO term. Using standard GO term lists such as ‘biological process’ or ‘cellular component’ will lead to enrichment of many non-specific GO terms, as the genes encode proteins that perform tasks in many different cell types and tissues. In our case, using uncurated lists may lead to organ non-specific enrichment, such as kidney-related or reproductive-related GO terms. However, since there are many known interactions between the brain and other systems (e.g. between blood circulation and migraine [43] or neural-immune interaction [44]) this may also be useful for discovering novel associations [45] or more general molecular processes.

For a first, more selective approach, GSEA allows the upload of any user-created GO term list. Therefore, it is recommended to carefully generate an experiment-specific GO term list containing only relevant GO terms to achieve a targeted and interpretable analysis.

### Summary

The present study compares the differential gene expression patterns of voxels with low and high fMRI activation probability in the mouse brain evoked by warm and hot thermal stimulation. Significant enrichment of gene ontology terms was found mainly in voxels with high activation probability - except for the dorsal striatum, where enrichment was found only in the voxels with low activation probability. Interestingly, GO terms associated with neuronal functions tended to be enriched in voxels with high activation probability, above the expected percentage. Although based on gene expression data from male C57BL/6J mice housed at the Allen Institute for Brain Science, these publicly available gene expression data sets can be used to improve the interpretation of fMRI imaging data.

### Outlook

We plan to apply our method to different imaging modalities (e.g., PET [46]), fMRI techniques (e.g., rCBV [work in progress]), and stimulation methods (e.g., visual, mechanical, pinprick [work in progress]). We are next interested in using the GSEA pipeline to address differences in sex-or age-dependent activation patterns. We will also use the ABA connectivity data to investigate specific anatomical and functional connectivity differences between high and low AP voxels. We will also address the reverse question by asking which genes are most relevant for separating high from low AP voxels and whether there are specific gene expression patterns across brain regions. In addition, the ABA atlas system provides a cellular mouse brain atlas [47], paving the way for a new and more detailed level of investigation. In addition to the ABA, we aim to incorporate other (meta-) databases, including cell types (Allen Brain Cell Atlas [47] or [48]), imaging, connectivity and gene expression datasets [49], with the aim of extending our approach to other species, including humans [50].

Neuroscientists are attempting to refine the anatomical and functional parcellation of the brain to deepen our understanding of its underlying function through spatial-specific connectivity mechanisms. Recent studies have used resting state fMRI to generate more detailed, functionally-driven sub-parcellations of the (human) brain, particularly in the cerebral cortex [51–53]. In future projects, we aim to extend this work by integrating gene expression databases (e.g., ABA) with fMRI to add another dimension to functional parcellation. Specifically, we will perform correlation analyses between gene expression and (f)MRI-based functional and structural connectivity, as well as brain-wide analyses of the leading-edge gene sets of high AP voxels—gene expression that significantly discriminate voxels that respond to the stimulation from voxels that don’t.

## Methods

### Animals, fMRI procedures and calculation of fMRI mean activation probability maps

For this study, we used activation probability maps derived from 164 fMRI measurements, to represent a wide range of variability (101 maps from male C57BL/6N (39 individual mice), 43 from female C57BL/6N (18 individual mice) and 20 from female DBA1 (5 individual mice)). In short, mice underwent between one and seven sessions of thermal stimulation BOLD fMRI (gradient echo echo planar imaging: TR = 2000 ms; TEef = 25 ms; FOV = 15 mm * 15 mm; in-plane resolution 0.234 mm * 0.234 mm; slice thickness 0.5 mm; matrix 64 * 64 voxels; 22 coronal slices covering the brain from bregma-8.12 mm to 2.58 mm). During the fMRI session, three sets of ascending thermal stimuli were applied at the right hind paw of the mice (40 °C and 45 °C (innocuous, labelled ‘warm’); 50 °C and 54 °C (noxious, labelled ‘hot’)). The whole session lasted 50 min. For details about fMRI measurement and data processing see e.g. [39]. After standard general linear model analysis with combined predictors for ‘warm’ and ‘hot’ stimulation temperatures, the resulting statistical parametric maps were corrected for multiple comparisons using false discovery rate (FDR, q < 0.05). This led to binary activation maps containing the statistically significantly activated voxels per animal. Those binary activation maps were registered (as described in [39]) and averaged for warm and hot stimulation temperatures independently. Consequently, these maps represent the voxel-wise activation probability (1 = 100 % of the animals have a significantly activated voxel at this position).

Since our fMRI data have 22 slices and the ABA data have 66 slices in a different reference system, the average binary activation maps had to be registered into the ABA space.

First, we selected the 22 slices from the 200 µm ABA MRI template that best represented our anatomical grey-scale intensity-coded MRI slices. Second, we used the non-affine landmark registration module of 3D Slicer (v.4.8.1) to register our anatomical grey-scale data to this 22-slice-version of the ABA based on 88 manually selected landmarks using the thinplate registration method [42]. Third, we used the transforms module of v. 5.2.1 (v.4.8.1 has bugs in the transforms module but produces better results in creating the transforms based on the landmarks than v.5.2.1) to register the probability maps to the ABA. Finally, the 22 slices were upscaled to the 66 slices of the ABA using the resample scalar/vector/DWI volume module of v.5.2.1. Different software versions had to be used due to bugs in the respective modules.

### Preparation of the gene expression volume

The Allen Institute of Brain Science provides whole brain gene expression data from male C57BL/6J mice, comprising 19,422 genes, aligned in a 200 µm voxel grid [1, 2]. First, all available expression data sets that were not labelled as ‘failed quality control’ were downloaded from the website (25520 data sets) [2]. Data sets could be in coronal or sagittal sections, with the sagittal sections comprising only the left hemisphere. The data for the missing hemisphere were imputed by flipping the existing data and forming a maximum projection per voxel. It should be noted that this creates artificial bilaterality and that these data cannot be used to infer laterality effects.

If more than one data set existed per gene, a maximum projection was calculated of all the corresponding data sets. The final collective gene expression volume contained 19,422 genes, with one 3D expression data set per gene.

### Definition of the voxel number for GSEA

For our analyses, we wanted to compare the gene expression of voxels with the highest AP evoked by thermal stimulation with that of an equal number of voxels with the lowest AP. The AP range differed for all regions analyzed (cf. Fig 2). Ranging from 0%-56% for lowest (bottom) probability and 79%-89% for highest (top) AP, we decided to use a dynamic region-specific threshold of the 5% and 95% quantile (rounded) to identify the most and least active voxels determined by their AP. This resulted in a different number of voxels analyzed for each region (see S2 Table), but importantly, the same number of voxels for the bottom and top AP per brain region as required for GSEA analysis.

### Atlas Creation

The Allen Institute of Brain Science provides a 3D mouse brain atlas (Common Coordinate Framework v3 Atlas; CCF [19, 54]) which is spatially corresponding to their gene expression volumes. It contains > 900 annotated grey and white matter structures in multiple detail levels. For use with our fMRI data, such a high-resolution annotation is not feasible, as many regions would be too small. Therefore, we combined the ABA annotations to create our own ABA-derived brain atlas, containing 102 grey matter brain regions (204 for both hemispheres), 8 fiber tracts and the ventricle system (though the latter two were not used in this analysis). Keeping the ABA coordinate system allowed us to use our atlas variant in combination with data provided by the ABA (structural connectivity and gene expression data). Left and right brain regions were treated as one region due to the artificial bilaterality introduced (see above). We analyzed gene expression of thalamus (TH), somatosensory cortex (S1 and S2; CTXs), cingulate cortex (including retrosplenial; CG), insular cortex (INS), visual cortex (VIS), hippocampus (HC), dorsal (STRd) and ventral striatum (STRv), and cerebellum (CER). For proof of concept, we additionally analyzed motor-related pons (PONmr), piriform cortex (PIR), and hypothalamus (HYP).

### Calculation of voxel-wise expression data

As introduced by ABA, many gene expression data sets contain voxels where no gene expression was measured, (value-1), e.g. because brain sections had to be omitted due to quality issues. This must not be confused with voxels where the gene is not expressed (value 0). While the 0 voxels must be included in the analysis, the-1 voxels must be excluded. To address this, we first restricted the activation maps to the relevant atlas region and sorted the voxels by activation probability in descending order. For each gene, beginning with the voxel with highest activation probability, we checked if the expression value in the corresponding gene expression volume was greater than - 1. If so, the expression value was recorded for that gene. The algorithm continued down the voxel list, skipping any voxels with a value of-1, until valid expression data for 5% of the total voxels was obtained (see S2 Table). The same approach was used for the low-probability voxels, starting from the bottom of the activation probability list.

This process ensured that the analyzed voxels, although differing slightly between genes, always included the 5% with the highest or lowest activation probabilities that had measurable gene expression data. A spatial projection of the selected voxels is shown in Figure 1, and Supplementary Figure 2 provides a graphical representation of the workflow.

The algorithm also checked whether the voxel sets for high and low activation probabilities overlapped due to skipping too many voxels in a given gene expression found (which was rare, see S3 Table), it was recorded in a log file. This issue was particularly common in sagittal ABA gene expression datasets, where missing slices led to the absence of expression values for certain genes. This can introduce bias, especially in small brain regions, as skipping too many voxels may result in overlapping voxel sets for low and high AP, causing the exclusion of that gene from the analysis. In very small regions, this overlap can occur frequently. To ensure consistency, the same set of genes should be analyzed across all brain regions. However, including very small brain regions in the analysis may lead to the exclusion of many genes from all datasets, thus reducing the power of the GSEA analysis. Therefore, we restricted the analysis to brain regions with a total volume of at least > 500 voxels.

**S3 Table.**
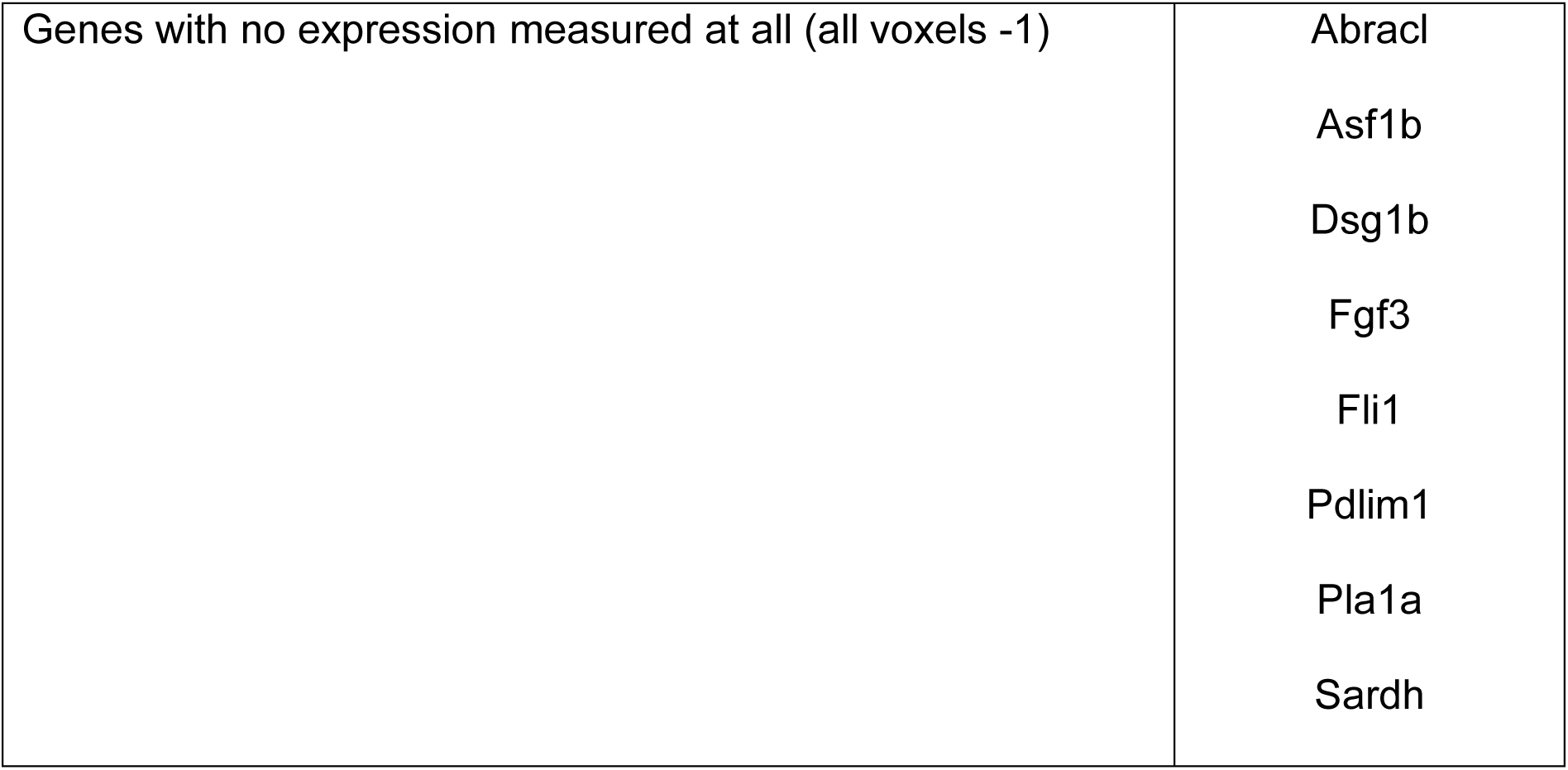
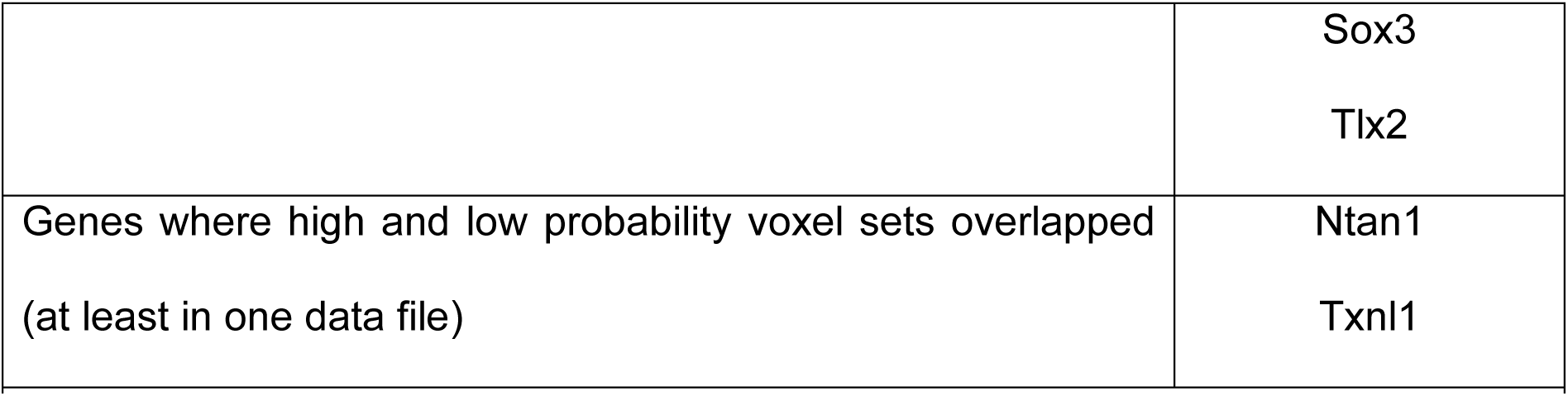
Genes removed from all data sets.

**S4 Table.**
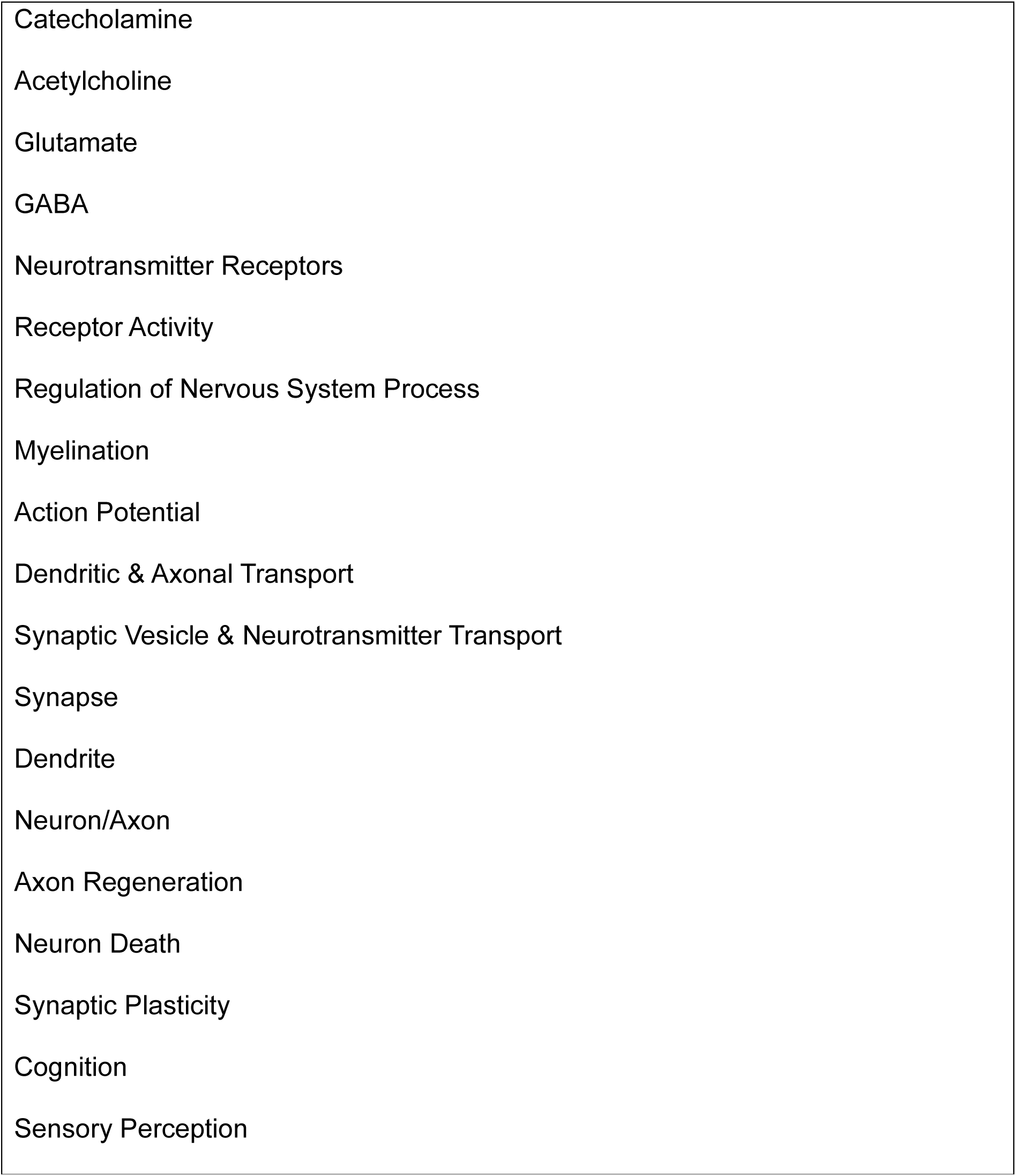
Super-GOs.

The algorithm recorded the gene names, the number of voxels (including the skipped voxels) needed to collect the 5% of voxels, log files, and expression values for the top and bottom 5% of voxels. One such data file was saved for each brain region, based on either warm or hot probability maps. To ensure comparability, we compiled all the data files at the end and removed any genes with missing or overlapping data from all datasets (for a list of deleted genes, see S3 Table). This resulted in a total number of 19,410 genes analyzed using GSEA.

### GSEA Analysis

The GSEA software (v. 4.3.2) from the Broad Institute was used for analysis [30, 55]. The resulting voxel-wise expression data for all 19,410 (after deletion of overlapping and missing genes) genes of all brain regions were arranged according to the GSEA guidelines (https://docs.gsea-msigdb.org/#GSEA/Data_Formats/), saved as txt-files with corresponding phenotype label files as cls-files.

GSEA was run for voxels with high AP versus voxels with low AP, for each stimulation temperature and each brain region independently. Mouse gene sets analyzed were m5.go.bp.v2023.2Mm.symbols.gmt (biological process, **BP**) and m5.go.cc.v2023.2Mm.symbols.gmt (cellular component, **CC**) provided by GSEA software. Additionally, we created curated versions of the **BP** and **CC** gene sets, containing only GO terms associated with neuronal and brain function (e.g. neurotransmitter secretion, memory, dopamine secretion, neuron remodeling, synapse, terminal bouton; termed ‘neuro-associated biological process’ (**naBP**) or ‘neuro-associated cellular component’ (**naCC**). The neuro-associated gene set **naBP** was further consolidated using the community detection algorithm ‘Enrichment Map’ [31] of Cytoscape 3.10.2 into 19 comprehensive Super-GOs for broader analysis (S4 Table). Analysis was run for each of these gene sets separately. For further details on GSEA Settings see S5 Table.

**S5 Table.** GSEA Settings.

| Fieldname | Parameter |
| --- | --- |
| Chip Platform | Mouse_Gene_Symbol_Remapping_MSigDB.v2023.2.Mm.chip |
| Permutation type | phenotype |
| Enrichment statistic | weighted |
| Metric for ranking genes | Signal2Noise |
| Gene list ordering mode | real |
| Max size | 500 (2000 for Super-GOs, as some of the merged data sets contained more than 500 genes) |
| Min size | 15 |
| Collapsing mode for probe sets=> 1 gene | Max_probe |
| Normalization mode | meandiv |
| Seed for permutation | timestamp |
| Randomization mode | no_balance |
| **S5 Table. GSEA Settings.** |  |

### Gene Ontology Term Enrichment Analysis

For each gene set (**BP**, **CC**, and Super-GOs), results were sorted according to their FDR q-value (p-value correction for multiple comparisons). The number of overall as enriched classified GO terms (independently for all FDR and FDR q < 0.2), significantly (FDR q < 0.05) enriched GO terms, and the number of significantly enriched GO terms with neuro-association (in case of **BP** and **CC**) were determined.

## Supporting Information

**S1 Fig. Spatial distribution of activation probability.** Shown are representative slices of averaged FDR-corrected activation maps from 164 warm and hot thermal stimulation fMRI measurements. The color code indicates the percentage of measurements in which each respective voxel was significantly activated by the stimulation.

(PDF)

**S2 Fig. Schematic representation of our GSEA-workflow.**

(PDF)

**S3 Fig. Number of enriched GO terms in the top and bottom 5% voxels.** Shown are all enriched GO terms (grey), GO terms with FDR < 0.2 (commonly used for gene expression analysis; light blue/red), and GO terms with FDR < 0.05 (usually used in imaging data; blue/red). From the latter ones, the subpopulation of GO terms associated with neurons, neurobiological processes or localizations are marked in darker color.

(PDF)

**S4 Fig. Span in mean AP does not correlate with significant GO term enrichment.**

(A) Frequency of AP. The black bars represent a histogram of the frequency of the AP (x-axis) across all voxels of the brain region. Blue and red dots represent the average AP of the top and bottom 5% voxels analyzed for each gene.

(B) Spatial distribution of the 5% analyzed voxels.

(C) Enriched GO terms. Bar plots show all enriched GO terms (grey), GO terms with FDR < 0.2 (commonly used for gene expression analysis; light red), and GO terms with FDR < 0.05 (usually used in imaging data; red). From the latter ones, the subpopulation of GO terms associated with neurons, neurobiological processes or localizations are marked in dark red.

(PDF)

**S5 Fig. Enrichment of the summarized Super-GO terms for all voxels using the manually curated neuro-associated BP gene list.**

Bubble size represents the size of the summarized gene set, color codes the FDR q-value and x-axis shows the normalized enrichment score (NES).

(PDF)

**S6 Fig. Volumetric representation of sensory, association, and visual cortex.** High AP voxels of VIS (red voxels) are located at the border to sensory (light yellow voxels) and association cortex (light green voxels), while low AP voxels (blue voxels) are more concentrated in the core of VIS.

(PDF)

**S7 Fig. GSEA analysis of 5% random voxels.** Spatial projection of the 5% random voxels (A). Number of enriched GO terms for the gene set **BP** (B).

(PDF)

## Acknowledgements

We’d like to thank the Allen Institute for Brain Science for making their data publicly available (from https://mouse.brain-map.org/).

AH reports funding from DFG CFU 5024 GB.Com. IW reports funding from IZKF Erlangen ELAN P139.

None of the funding sources had any role in the conception of the study, the development of the methods, the analysis or interpretation of the data, the writing of the manuscript, or the decision to submit the paper for publication.

## Author Contributions

IW: analysis, development of methodology and software, data validation, visualization and writing of the original draft; AA: software; SK: software and data validation; MH: data validation; AH: supervision and conceptualization, development of methodology and writing of the original draft.

All authors discussed the results, reviewed, edited and commented on the initial draft.

## Data availability statement

Gene expression data used in this study is publicly available from https://mouse.brain-map.org/.

FMRI data will be available upon request.

