## Supplementary Figures PDF for "Gene expression patterns decompose fMRI activation in a sub-region-specific manner in mice after nociceptive stimulation"

### - Supplementary Material -

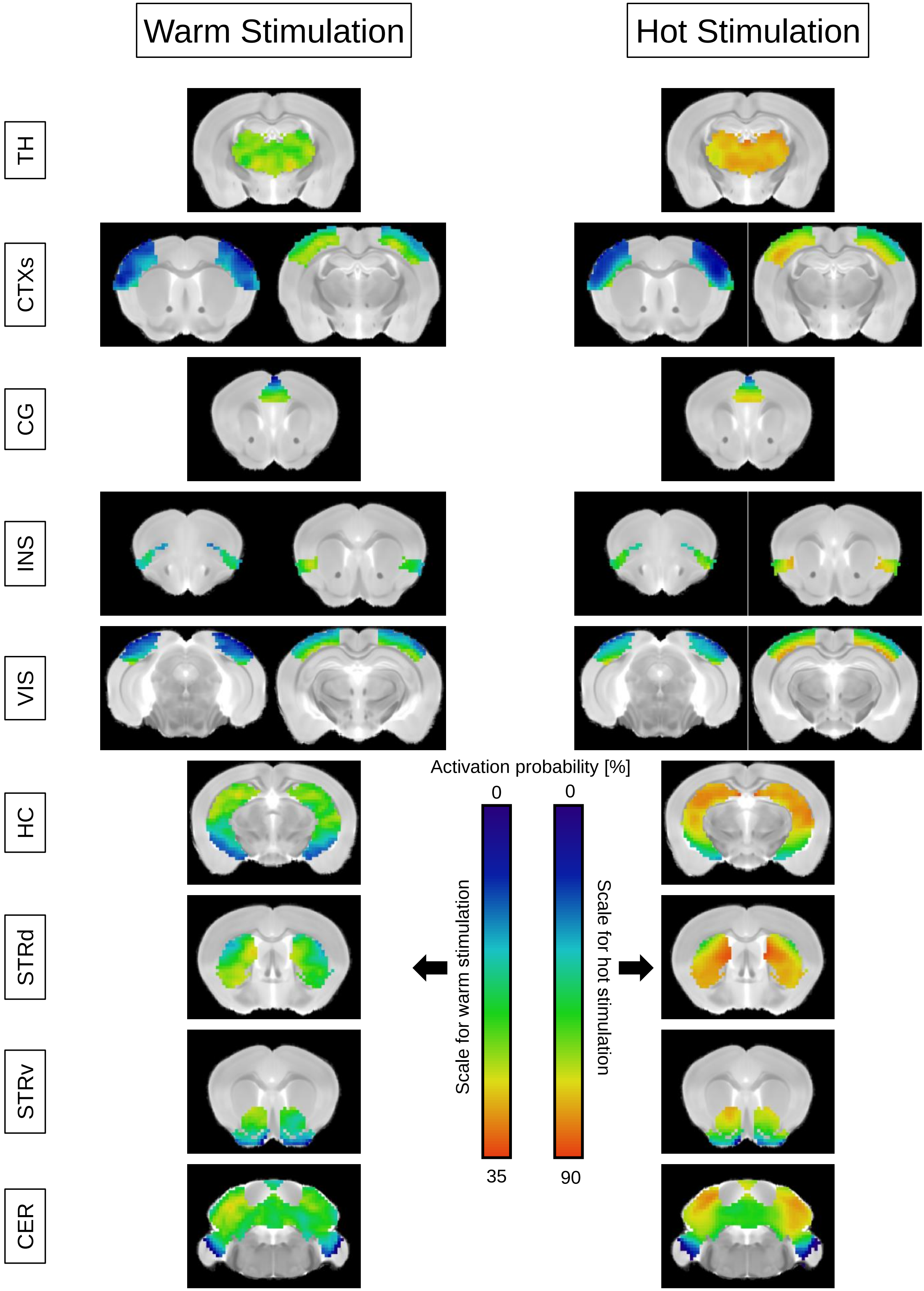

Warm Stimulation
Hot Stimulation
TH
CTXs
CG
INS
VIS
HC
STRd
STRv
CER
Activation probability [%]
0
0
Scale for warm stimulation
Scale for hot stimulation
35
90

**S1 Fig. Spatial distribution of activation probability.**

Shown are representative slices of averaged FDR-corrected activation maps from 164 warm and hot thermal stimulation fMRI measurements. The color code indicates the percentage of measurements in which each respective voxel was significantly activated by the stimulation.

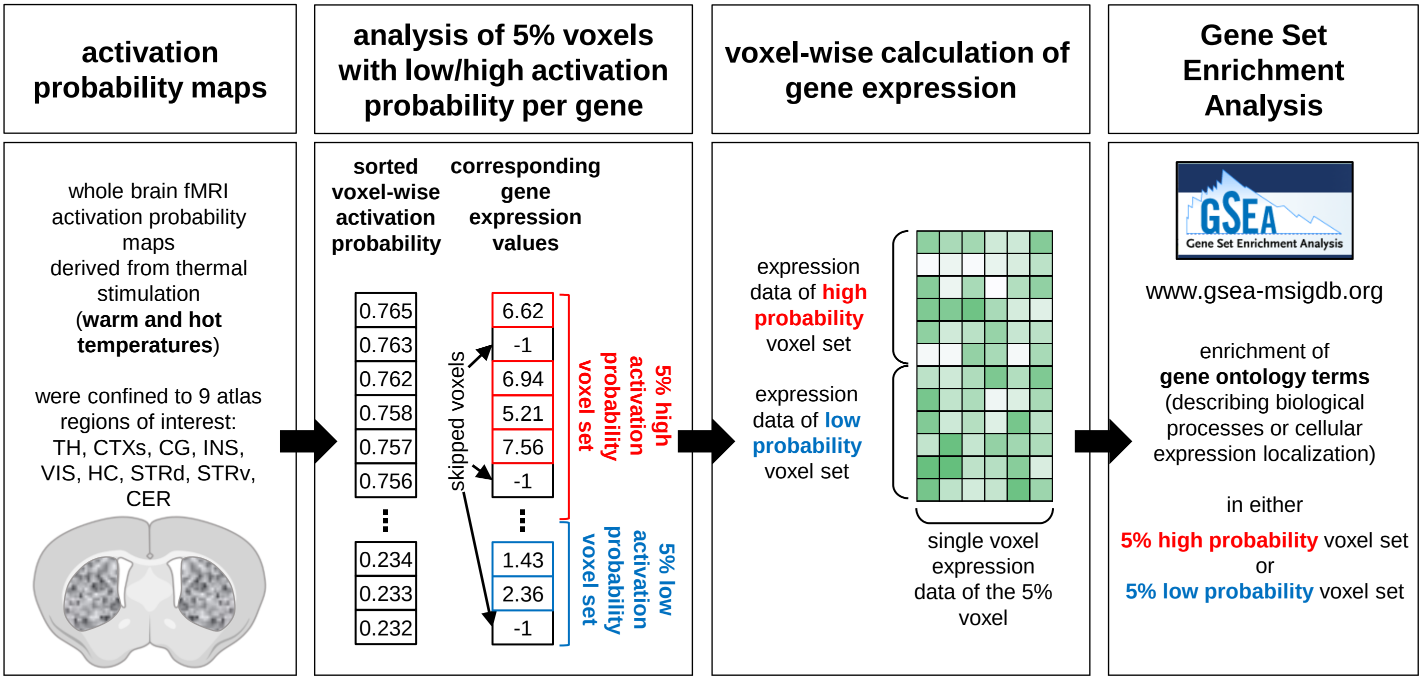


**S2 Fig. Schematic representation of our GSEA-workflow.**

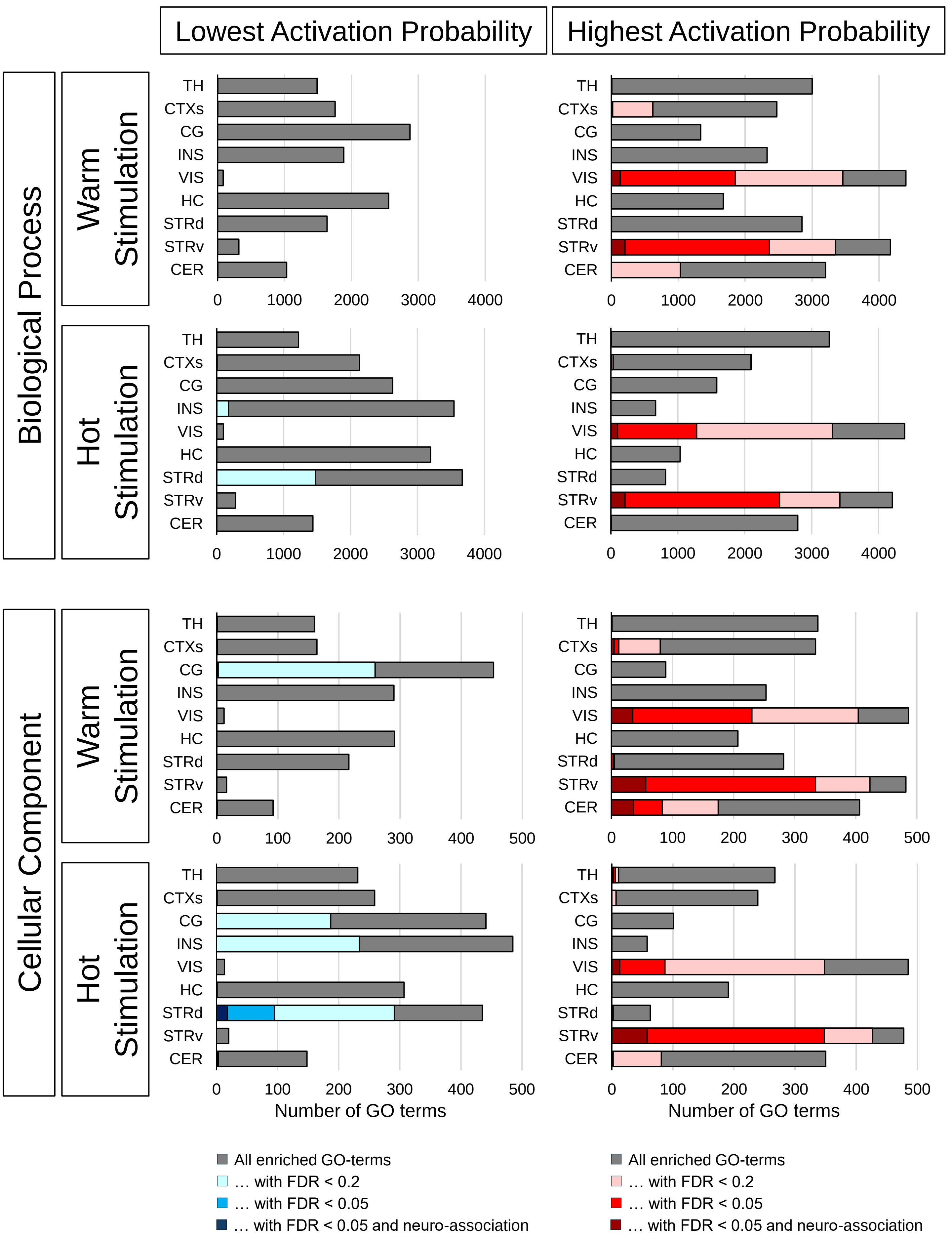
Lowest Activation Probability
Highest Activation Probability
Biological Process
Cellular Component
Warm Stimulation
Hot Stimulation
TH
CTXs
CG
INS
VIS
HC
STRd
STRv
CER
0
1000
2000
3000
4000
100
200
300
400
500
Number of GO terms
All enriched GO-terms
… with FDR < 0.2
… with FDR < 0.05
… with FDR < 0.05 and neuro-association

**S3 Fig. Number of enriched GO terms in the top and bottom 5% voxels.**

Shown are all enriched GO terms (grey), GO terms with FDR < 0.2 (commonly used for gene expression analysis; light blue/red), and GO terms with FDR < 0.05 (usually used in imaging data; blue/red). From the latter ones, the subpopulation of GO terms associated with neurons, neurobiological processes or localizations are marked in darker color.

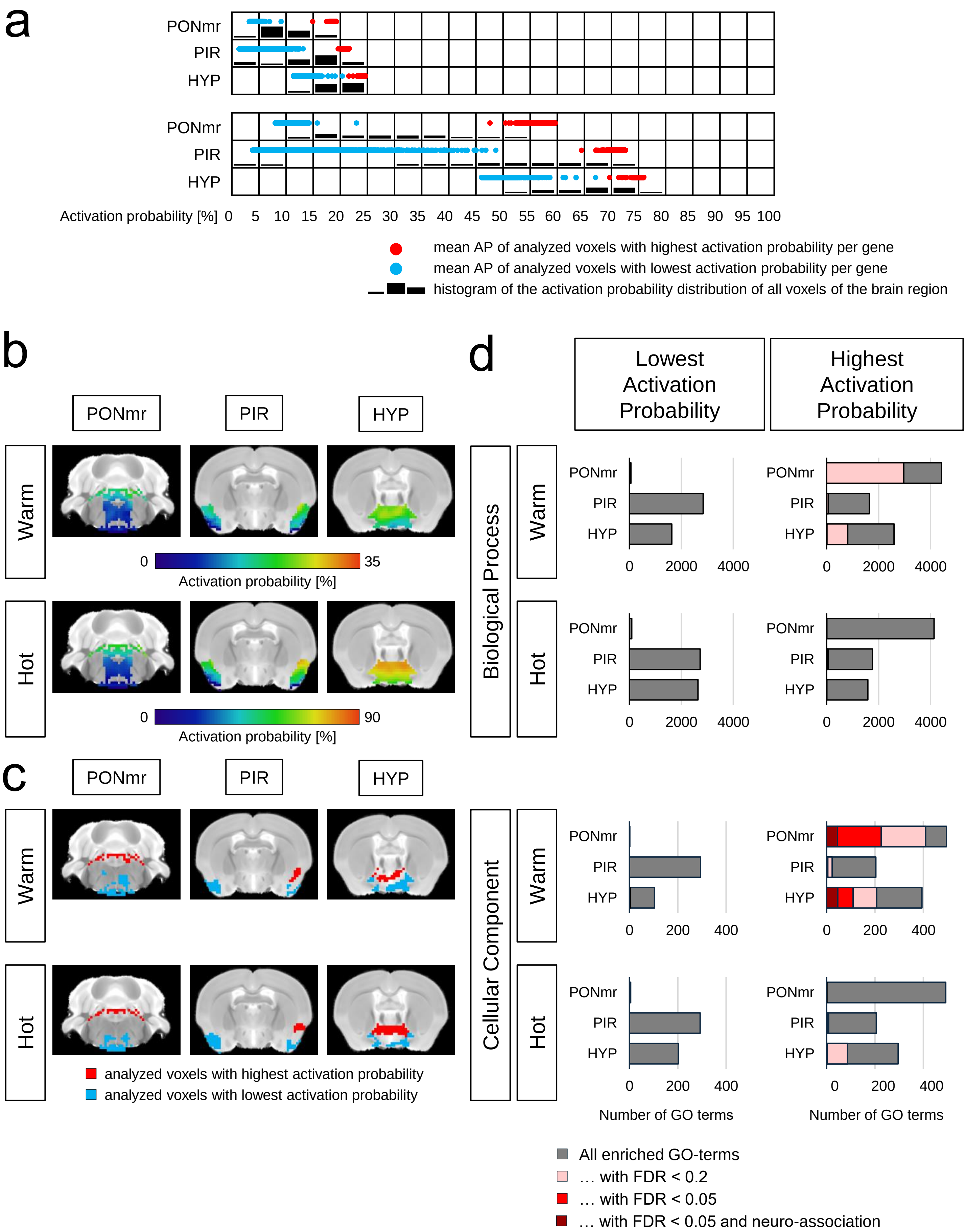

a
PONmr
PIR
HYP
Activation probability [%] 0 5 10 15 20 25 30 35 40 45 50 55 60 65 70 75 80 85 90 95 100
mean AP of analyzed voxels with highest activation probability per gene
mean AP of analyzed voxels with lowest activation probability per gene
histogram of the activation probability distribution of all voxels of the brain region
b
Warm
Hot
0 35
Activation probability [%]
0 90
c
analyzed voxels with highest activation probability
analyzed voxels with lowest activation probability
d
Lowest Activation Probability
Highest Activation Probability
Biological Process
Cellular Component
0 2000 4000
0 200 400
Number of GO terms
All enriched GO-terms
… with FDR < 0.2
… with FDR < 0.05
… with FDR < 0.05 and neuro-association

**S4 Fig. Span in mean AP does not correlate with significant GO term enrichment**.

(A) Frequency of AP. The black bars represent a histogram of the frequency of the AP (x-axis) across all voxels of the brain region. Blue and red dots represent the average AP of the top and bottom 5% voxels analyzed voxels for each gene.

(B) Spatial distribution of activation probability.

(C) Spatial distribution of the 5% analyzed voxels.

(D) Enriched GO terms. Bar plots show all enriched GO terms (grey), GO terms with FDR < 0.2 (commonly used for gene expression analysis; light red), and GO terms with FDR < 0.05 (usually used in imaging data; red).

From the latter ones, the subpopulation of GO terms associated with neurons, neurobiological processes or localizations are marked in dark red.

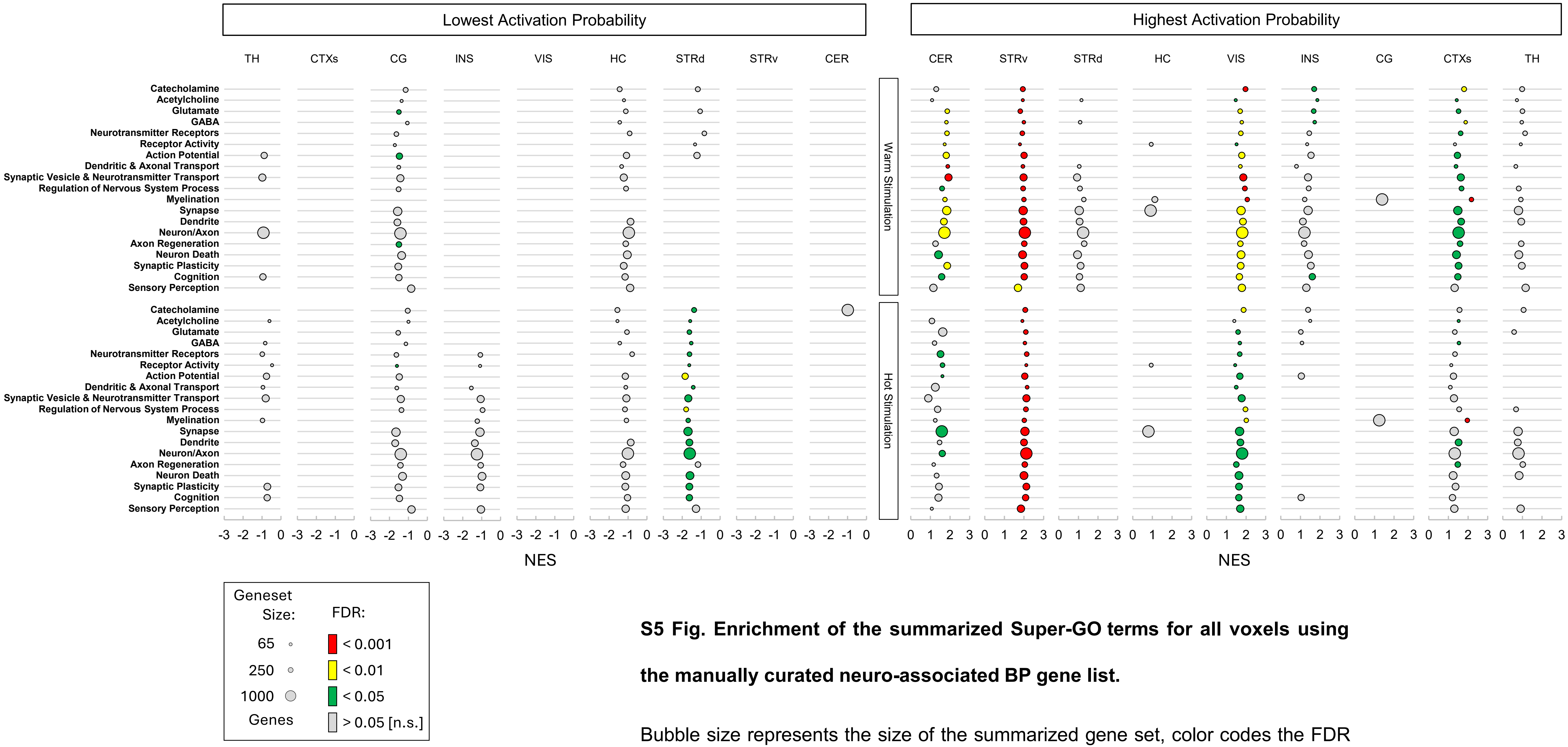


**S5 Fig. Enrichment of the summarized Super-GO terms for all voxels using the manually curated neuro-associated BP gene list.**

Bubble size represents the size of the summarized gene set, color codes the FDR q-value and x-axis shows the normalized enrichment score (NES).

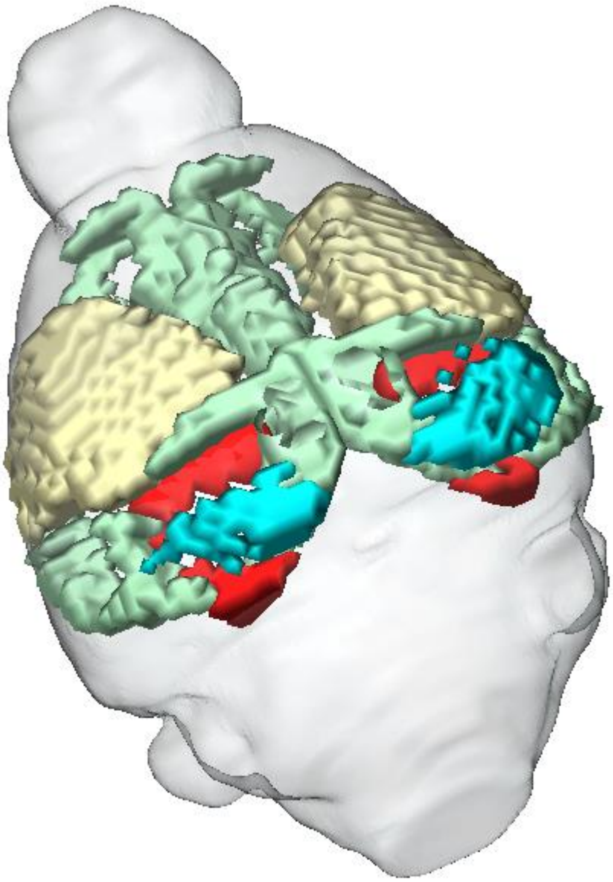

VIS analyzed voxels with highest activation probability
VIS analyzed voxels with lowest activation probability
somatosensory cortex area
association cortex area

**S6 Fig. Volumetric representation of sensory, association, and visual cortex.** High AP voxels of VIS (red voxels) are located at the border to sensory (light yellow voxels) and association cortex (light green voxels), while low AP voxels (blue voxels) are more concentrated in the core of VIS.

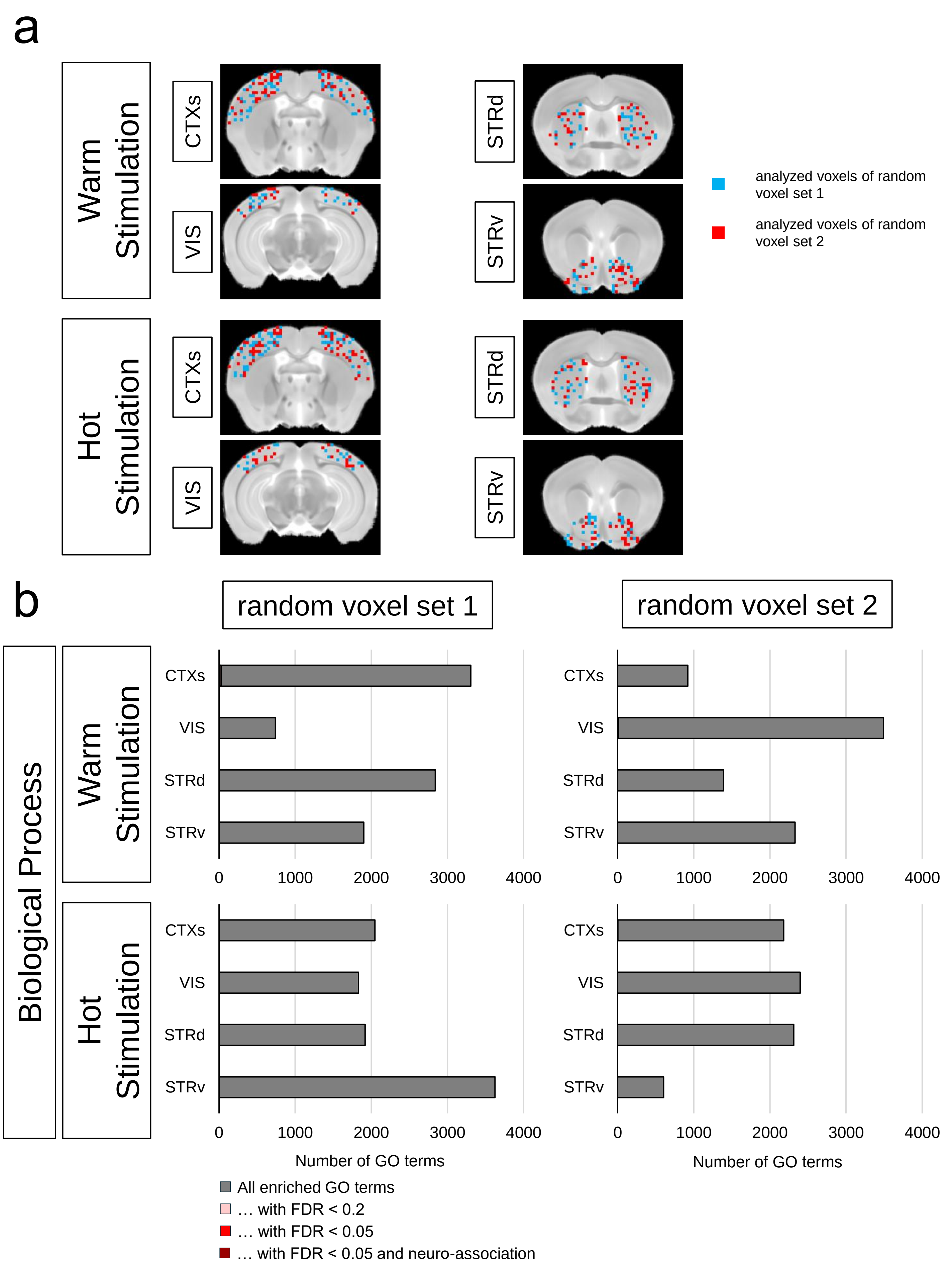
a
Warm Stimulation
Hot Stimulation
CTXs
VIS
STRd
STRv
analyzed voxels of random voxel set 1
analyzed voxels of random voxel set 2
b
random voxel set 1
random voxel set 2
Biological Process
0
1000
2000
3000
4000
Number of GO terms
All enriched GO terms
… with FDR < 0.2
… with FDR < 0.05
… with FDR < 0.05 and neuro-association

**S7 Fig. GSEA analysis of 5% random voxels.** Spatial projection of the 5% random voxels (A). Number of enriched GO terms for the gene set BP (B).
